# A single protein to multiple peptides: Investigation of protein-peptide relationship using targeted alpha-2-macroglobulin analysis

**DOI:** 10.1101/2022.12.11.519938

**Authors:** Pelin Yildiz, Sureyya Ozcan

**Affiliations:** Department of Chemistry, Middle East Technical University, 06800 Ankara, Turkiye; Nanografi Nanotechnology Co, Middle East Technical University, Technopolis, 06531 Ankara, Turkiye; Cancer Systems Biology Laboratory (CanSyL), Middle East Technical University, 06800 Ankara, Turkiye

**Author notes:** **Corresponding Author Sureyya Ozcan** – Department of Chemistry, Middle East Technical University, 06800 Ankara, Turkey.

**Keywords:** Targeted proteomics, mass spectrometry, quantitative proteomics, alpha-2-macroglobulin, unique peptide, serum

## Abstract

Recent advances in proteomics technologies have enabled analysis of thousands of proteins in a high-throughput manner. Mass Spectrometry (MS) based proteomics, uses a peptide centric approach where biological samples undergo a specific proteolytic digestion and then only unique peptides are used for protein identification and quantification. Considering the fact that a single protein may have multiple unique peptides and a number of different forms, it becomes essential to understand dynamic protein-peptide relationship to ensure robust and reliable peptide centric protein analysis. In this study, we investigated the correlation between protein concentration and corresponding unique peptide responses under conventional proteolytic digestion conditions. Protein-peptide correlation, digestion efficiency, matrix-effect, and concentration-effect were evaluated. Twelve unique alpha-2-macroglobulin (A2MG) peptides were monitored using a targeted MS approach to acquire insights into protein-peptide dynamics. Although the peptide responses were reproducible between replicates, protein-peptide correlation was moderate in protein standards and low in complex matrices. The results suggest that reproducible peptide signal could be misleading in clinical studies and a peptide selection could dramatically change the outcome at protein level. This is the first study investigating quantitative protein-peptide correlations in biological samples using all unique peptides representing the same protein and opens a discussion on peptide-based proteomics.

## INTRODUCTION

Clinical proteomics facilitates the analysis of large number of proteins and aims to discover new biomarkers and novel targets^1,2^. In bottom-up (shotgun) proteomics, peptide-centric approach is used as first proteins are digested using proteases, then peptide products are analyzed by mass spectrometry (MS)^3^. The method focuses on identification and/or quantification of unique peptide(s) of the protein, also known as proteotypic or signature peptide(s), to ensure specificity of the analysis. Considering the protein size and sequences proteins often contain multiple unique peptides. Since it may not be feasible to use all the unique peptides for each target protein, one or a few peptides often used in practice for years for targeted proteomics studies^4–8^. The measures used to select a peptide of interest applied in common practice are i) the peptide length (8-25 amino acids), ii) lack of post-translational modifications (PTMs; glycosylation, phosphorylation, etc), iii) avoidance of chemically active amino acids (methionine (M), cysteine (C), glutamine (Q), tryptophan (W), histidine (H), and asparagine (N)) residues that can cause oxidation, acetylation and iv) no mis-cleavages^9–13^. In practice, there is a wide variety of tools available for peptide selection^14–22^. It is adopted that all unique peptides in the protein should behave the same. Therefore, the tendency is to perform either a literature mining and select the target or choosing the most intense peak after an MS scan for the analysis. Thus, the peptide representing the same target protein often varies from one study to another.

The tryptic digestion has been extensively used in clinical proteomics for various biological samples including serum, plasma, tissue and cell^23,24^. Several processing conditions are used in the literature when the biological samples are prepared using various methods^25–34^. Trypsin is accepted as ‘the gold standard enzyme’ for protein digestion and process involves protein denaturation, reduction and alkylation prior to enzymatic digestion^23,35^. Although the ultimate aim is to facilitate efficient digestion by producing specific peptides and avoid mis-cleavages, there is often not enough quality control measures applied in the studies to control the processing^36,37^. Additionally, many of the reagents used in protein preparation limit trypsin activities, and highly folded proteins are resistant to proteolysis^38^. Lysine (K) amino acid residue cleavage is particularly difficult during trypsin digestion^39,40^. These limitations are improved when trypsin and Lys-C are combined^38,41^. The studies’ results demonstrate that the combination of trypsin and Lys-C enzymes boosted the number of tryptic peptides found throughout the digestion process by minimizing mis-cleavage^38,41,42^.

There are huge efforts to standardize and validate analytical protocols for clinical proteomics^4,8,20,43^. Lee et al. originally introduced the concept of the “fit-for-purpose” approach for biomarker method validation in the early 2000s. This concept means that assessments need to be validated in consideration of the intended use of the data and any applicable regulatory requirements. Any further validation might be required iteratively when the data’s intended purpose changes^44,45^. The basis of potential targets that can be evaluated in the early phases of biomarker development is intended to be as broad as possible, and when biomarkers advance toward clinical sufficiency, analytical performance is designed to be as high as possible. According to the approach, the validation of targeted proteomic assays based on Multiple Reaction Monitoring (MRM) is also subject to the typical characteristics of analysis such as linearity, detection limit, repeatability, selectivity, and stability. Although the results of the assay are highly reproducible at the protein and peptide levels in accordance with the “fit-for-purpose” approach in plasma and tissue samples^8,46^, the results are generally restricted with a single peptide^8,46^. For instance The Clinical Proteomic Tumor Analysis Consortium (CPTAC) assessed the reproducibility, recovery, linear dynamic range and limits of detection and quantification of 11 peptides representing 7 proteins using an MRM-based protein quantification at 8 different sites^4^. The results indicate a good precision and reproducibility between the laboratories. Nevertheless, multiple peptides representing the same protein were not correlated to each other. The differences in the peptide intensities were reported as the variations in the ionization efficiencies. The key question of why and on what basis specific numbers of unique peptides are selected to represent proteins remains unresolved, even in fit-for-purpose approaches^4,8,43,46^. This demonstrates how crucial it is to choose unique peptides for the representation of proteins.

In addition, studies have shown that optimizing the digestion conditions could effectively reduce the nonspecific peptide cleavages^47^. Similarly, there are different approaches applicable to improve protein extraction as well as peptide ionization^48^. Despite the fact that trypsin is a highly specific protease, a large number of non-specific tryptic peptides are widely known with certainty. Discovering peptides originating from non-tryptic pathways might help to improve sequence coverage, however screening for all possible nonspecific peptides in shotgun proteomics data from complicated samples such as plasma, serum, and saliva can be difficult^49^. The amount of non-specific trypsin cleavages within a variety of biological systems should be investigated, as well as the differences in non-specific trypsin cleavage incidence among different sample matrices. Major experimental parameters such as protein:enzyme ratio, incubation time and temperature should be optimized and, if it is necessary, modified during sample preparation to control non-specific trypsin cleavages. In addition to the variation both protein and peptide level that could cause from the analysis parameters, the extend of the biological variation is unknown. The size of the proteome and protein forms in a single system is still a big debate^50^. Plubell *et al*. recently challenged the protein quantification and stated that as different protein form may yield different peptides, peptide centric approach may cover the biological differences^9^. The issue is addressed in the article as follows: “*Fundamentally all bottom-up acquisition types are all subject to the same problem. The main difference will be in the number of peptides being measured, which will influence the possible interpretation of results. As we stated in the manuscript; “*…*if coverage is low, differences in peptides specific to a proteoform may not be captured*.*” Therefore, if we choose to only measure 2 or 3 peptides per protein in a targeted assay, we may miss out on detecting interesting or relevant differences in other peptides*.*”* Peptides, reported in the literature, on which the result of protein measurement is based might be an incomplete picture as all other peptides may not be correlated to each other. To investigate the protein-peptide and peptide-protein relation we used a model protein. Alpha-2-macroglobulin (A2MG), is a plasma glycoprotein that is present in a number of biological fluids such as blood, serum, and saliva^51,52^. Due to its ability to attach to foreign peptides and molecules, it is biologically active because it inhibits a variety of proteases and functions as a disease protection barrier. A2MG’s multiple reactive sites enable it to assist in the reversible or irreversible capture of proteins with a variety of biological activities^53^. A2MG has established into a biomarker for a wide range of diseases. A2MG a clinically important protein associated with liver^54^, diabetes^55^, neurological diseases^56–58^, and prostate cancer^55^, was selected as a reference protein. Recent structural investigations of the A2MG protein in the literature have mostly focused on the protein’s conformational changes and complex formation^59–61^, binding interaction with drugs/small substrates^62–69^, and PTMs^70^. The position of the unique peptide in the protein structure and commonly observed mis-cleavages were examined in this study ^71,72^.

The dynamic protein-unique peptide interaction is yet unknown, despite the widespread belief that all unique peptides that reflect the protein of interest behave in the same manner. The aim of this study is to examine the dynamic quantitative protein-unique peptide relationship beyond ionization efficiencies. To this end, A2MG was selected as a representative protein and serum was employed as a model system. The proteolytic digestion was applied to various concentrations of samples and all unique peptides representing A2MG were monitored. The correlation of protein and peptide was examined through a targeted MRM method. The outcomes of this research have potential to be utilized for clinical applications towards effective and reproducible biomarker research.

## MATERIALS AND METHODS

### Reagents

The peptides and internal standard were synthesized by PeptiTeam Ankara, Turkey. The following chemicals were used: A2MG protein standard (Sigma-Aldrich), fetal bovine serum (Sigma-Aldrich), human serum (Sigma-Aldrich), trypsin (Promega), trypsin/Lys-C mixture (Promega), ammonium bicarbonate (ABC, Sigma-Aldrich), dithiothreitol (DTT, Sigma-Aldrich), iodoacetamide (IAA, Sigma-Aldrich), LC-MS grade formic acid (Merck), LC-MS grade water (Merck), and LC-MS grade acetonitrile (Merck).

### Samples

The similarities and differences in the behavior of unique peptides in biological systems were investigated using reference systems; peptide standards (FEVQVTVPK, QGIPFFGQVR, VGFYESDVMGR, DMYSFLEDMGLK, LPPNVVEESAR), the A2MG protein standard, bovine and human serum. The peptide and protein standards were used for MS performance evaluation and analytical method optimization. Two levels of quality controls (QCs) were used to check the instrument’s stability and sample processing variation. Method performance was controlled to prevent systematic error. QCs were made for pooled human serum digest and a mixture of peptide standard samples.

### In-Solution Protein Digestion

Both trypsin and trypsin/Lys-C mixture digested triplicates were treated under the same circumstances for the initial reproducibility analysis. 5 μl thawed aliquots of serum and protein standard were digested by 10 μl of 45 mM DTT in 50 mM ABC at 65°C for 45 minutes, 10 μl of 100 mM IAA in 50 mM ABC in the dark for 30 minutes at room temperature, trypsin at a protein:enzyme ratio of 50:1, and trypsin/Lys-C enzyme mixture at a protein:enzyme ratio of 50:1. 50 mM ABC buffer solution was added to each triplicate so that the final total volume was same. All prepared triplicates were vortexed before being incubated at 37°C for 16 hours. After the incubation, a 1% (v/v) stock formic acid solution was added to the triplicates at a final concentration of 0.1% (v/v) to stop the digestion. During the experiment, fresh stock solutions were prepared. The internal standard was added to all prepared samples before the mass spectrometric analysis.

### LC-MS Analysis

Quantitation of all samples was performed by an Agilent 1260 Infinity II HPLC system coupled to an Agilent 6470A triple-quadrupole (QQQ) system (Santa Clara, CA). Agilent MassHunter Quantitative Analysis software was used for data acquisition and processing. Each sample was analyzed using LC–MS in triplicate. Chromatography was performed with mobile phase A LC-MS grade water containing 0.1% formic acid and mobile phase B pure acetonitrile containing 0.1% formic acid. Peptides were eluted flow rate at 0.3 ml/min for 15% of mobile phase B for 4 minutes, increased to 65% and held at 65% mobile phase B for 2.4 minutes before returning to 5% mobile phase B. The Agilent InfinityLab Poroshell 120 C18 (3.0 × 150 mm, 2.7 μm) column was used as an HPLC column. The reversed-phase C18 column was incubated during analysis in an oven at 50°C for better separation. The peptides were introduced to the QQQ by electrospray ionization (ESI) with Jet Stream of Agilent Technologies after being separated in the LC system. In the quadrupole 1 (Q1) and quadrupole 3 (Q3) compartments, each scan window was operated at a unit resolution. In the analysis, the dynamic MRM mode was selected with retention time windows of 2 minutes. The MS operating parameters were acquired with gas temperature of 300°C, gas flow of 11 L/min, nebulizer pressure of 40 psi, sheath gas temperature of 400°C, sheath gas flow of 11 L/min, capillary voltage of 3500 V, collision energy between 6.9 – 22.4 V.

### Data Analysis

Raw MS data were taken from the Agilent QQQ instrument in .d files format. The results were loaded into the Skyline software package (version 20.2.0.343, MacCoss Lab, UW). Data pre-processing was done manually by using Skyline software. The pre-processed MS data was subjected to statistical data analysis. Briefly, the MS data were normalized, correlated, and then, distribution and clustering analyses were done, respectively. Statistical data were treated and visualized by using OriginPro 2018 SR1 software (version b9.5.1.195, OriginLab Corporation).

## RESULTS AND DISCUSSION

We used a systematic approach focusing on a single protein and its multiple unique peptides to understand the quantitative connection. First, all unique peptides representing A2MG were identified. The targeted MS method was then optimized using a standard protein. Varying concentrations of A2MG protein was spiked on buffer, human serum and bovine serum to assess the matrix effect. All samples were digested using a conventional method using two common proteases; trypsin and trypsin/Lys-C mixture prior to MS analysis. The protein-peptide correlations were extensively studied to observe the dynamic changes that occur on protein level.

### The Selection of Unique Peptides of A2MG Protein

In order to investigate the dynamic unique peptide-protein relationship, A2MG was used as a reference protein. First, the A2MG protein sequence was obtained from the UniProt^73^ database, and then tryptic peptides were calculated using the ExPasy PeptideCutter^74^ interface. As a result of this analysis, 124 tryptic peptides of protein were obtained. Supplementary **Figure S1** shows the enzyme cleavage sites. The lysine (K) and arginine (R) sites in the A2MG protein’s 3D structure are highlighted in purple and yellow, respectively. 80 of the 124 tryptic peptides were cleaved from the lysine (K) residues, while 44 of them were cleaved from the arginine (R) residues. The uniqueness of the peptides was determined using three independent databases: UniProt, NextProt^75,76^, and BLAST^77^. The outcomes of the search were compared to ensure protein specificity. At the end of this analysis, 15 unique peptides were determined among 124 tryptic peptides. These unique peptide compositions, their positions, and sizes are shown in **Table S1**. The number of amino acids they contain varies between 8-30. Fifteen unique peptides were screened using an LC-MS and peptides were further filtered, and three (ALLAYAFALAGNQDK, GGVEDEVTLSAYITIALLEIPLTVTHPVVR, and QTVSWAVTPK) were excluded from the study due to low ionization efficiency. Peptide length influences mass spectrometry-based sequence identification. The optimum peptide length range to be studied for unique peptide is given in the literature as 8-25 amino acids^78^. Peptides, including six amino acids or less^23^, are small peptides, as they are not specific peptides. The peptide GGVEDEVTLSAYITIALLEIPLTVTHPVVR is also not in the optimal peptide length range. It was excluded from the study because too-long peptides interfere with the enzyme’s digestion ability, resulting in incomplete enzymatic digestion. On the other hand, because of a surrogate matrix, the unique peptide QTVSWAVTPK was also not considered in the study as a unique representative peptide. It is not just a unique peptide for humans. Using the UniProt BLAST software, it is detected in other mammals such as cattle, Sumatran orangutans, brown rats, and house mice. In addition to this information, the criteria used to select unique peptides are summarized in **Table S2** as peptide length, peptide active amino acid residues, and enzyme efficiency. Each unique peptide is colored red in the table to indicate the number of chemically active amino acid residues. We further investigated the behavior of these 12 unique peptides and their correlations under different biological conditions.

### The Localization of Unique Peptides

The locations of unique peptides in the 3D structure of A2MG protein (PDB:4ACQ^56^) was visualized in PyMOL^79^ in **Figure 1**. While **Figure 1A** shows the distribution of unique peptides on the protein structure, **Figure 1B** includes glycosylation sites to the unique peptide distribution. Each unique peptide is represented in **Figure 1A** by a different color, and the dark gray color represents the glycosylation sites in **Figure 1B** whereas the magenta color indicates all unique peptides. Furthermore, **Table S3** describes the color codes for unique peptides. The A2MG structure does not contain the bait region, between 690 and 728 amino acid sequences. Thus, the structural positions of unique peptides VGFYESDVMGR and LVHVEEPHTETVR could not be localized^80^.

**Figure 1.**
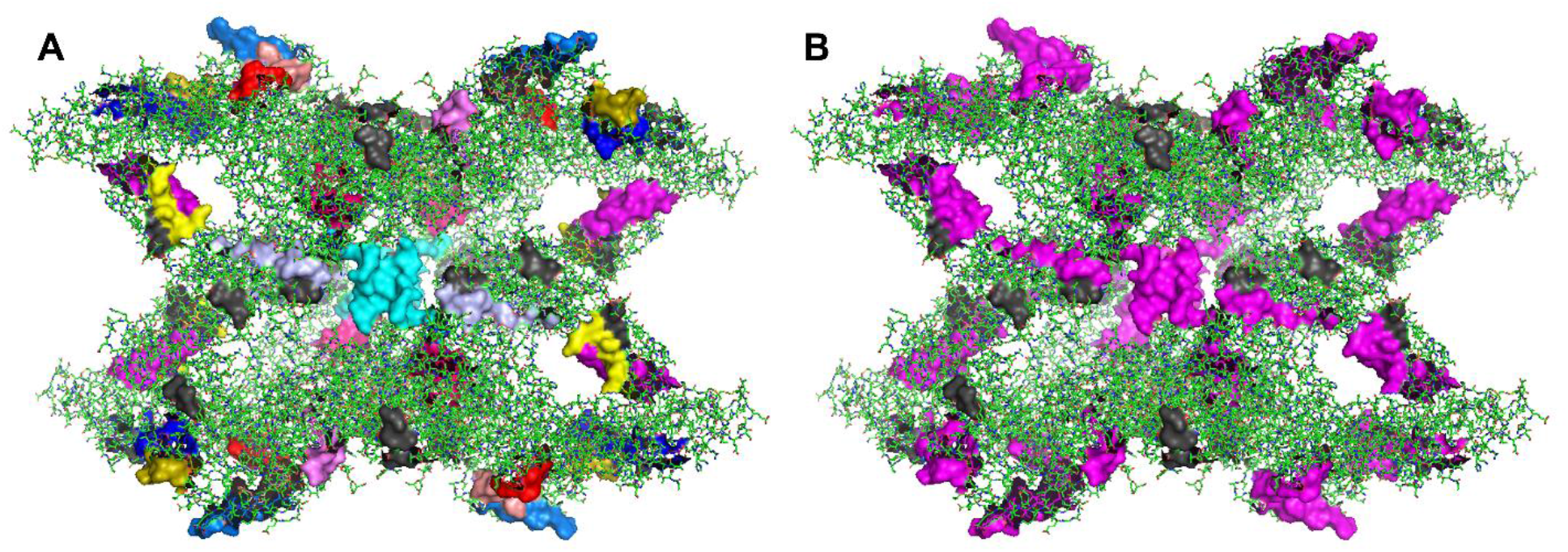
The 3D model of A2MG protein, shown as green sticks. **A**. The locations of unique peptides in the protein structure, shown as surface in different colors **B**. The locations of unique peptides (magenta) and glycosylation (dark gray) on the protein structure, shown as surface.

We can extrapolate from **Figure 1A** that the specific cleavage efficiency of unique peptides placed in the inner regions is predicted to be low, whereas the digestion efficiency of the peptides located in the outer region, which is easily accessible, is expected to be high. In addition, PTM is one of the important parameters when examining a protein’s structure. **Figure 1B** shows that the unique peptides are not located on the glycosylation sites. However, FEVQVTVPK, QGIPFFGQVR, and DTVIKPLLVEPEGLEK are positioned near to glycosylation sites, shown in **Table S4**, which could alter their digestion efficiency.

### The Association of Unique Peptides with Literature

After determining the locations of unique peptides on A2MG protein, the incidence of unique peptides used in studies published in the literature was analyzed. The frequency of unique peptides utilized in the literature for identification and/or quantification of A2MG protein is listed in **Table S5**. The relationship between the number of publications in this table and the unique peptides has been visualized using a word cloud graph to make it easier to interpret in **Table S5. Figure 2** shows that peptides with larger fonts are commonly investigated, whereas peptides with smaller fonts are rarely studied in the literature.

**Figure 2.**
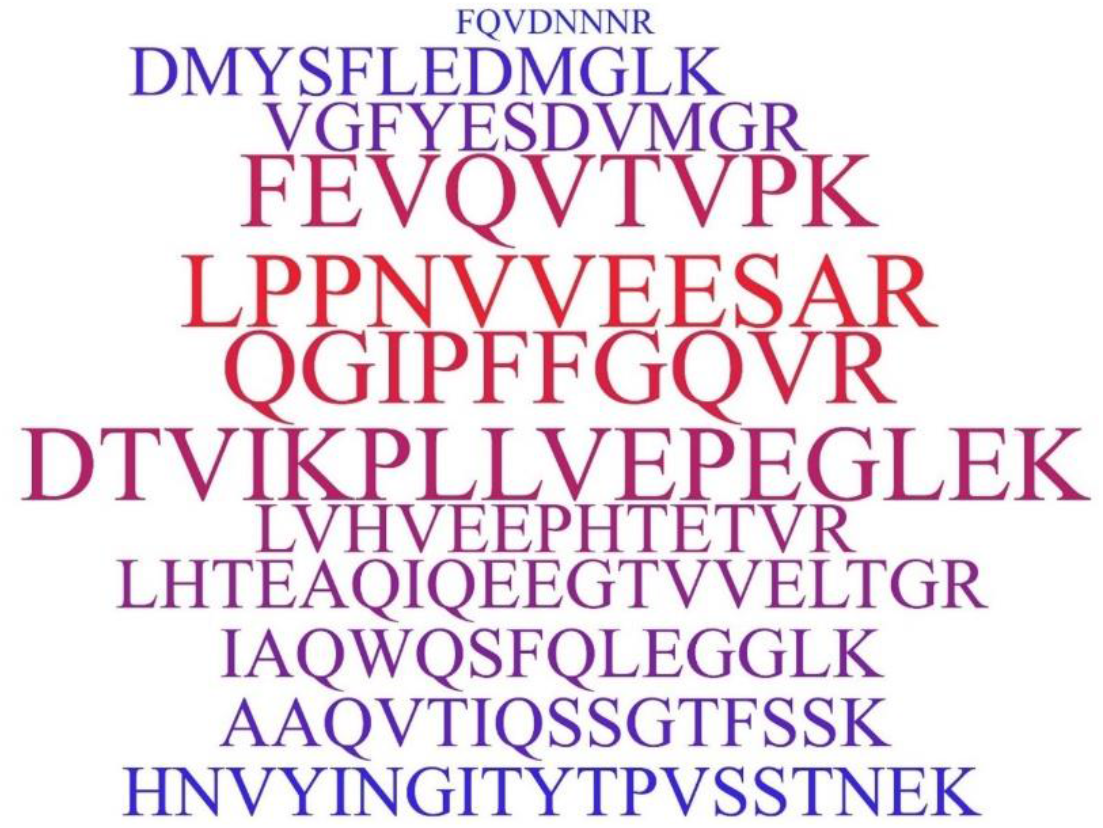
A word cloud plot represents the incidence of A2MG unique peptides in the literature.

The unique peptides, FEVQVTVPK, LPPNVVEESAR, QGIPFFGQVR, and DTVIKPLLVEPEGLEK, are frequently encountered. The peptide FQVDNNNR, on the other hand, has received the least amount of consideration in the literature. the four most studied unique peptides were found on the protein surface. The DTVIKPLLVEPEGLEK unique peptide containing mis-cleavage has been used for A2MG protein quantification in some studies^71,72^ Furthermore, the lengths of the frequently used unique peptides range from 9 to 16 amino acids. A unique peptide to represent a protein is usually chosen randomly or based on high abundance. Although there are no regulations regarding the selection of unique peptides to represent proteins, the literature has developed several standards. The following elements should be taken into account when choosing unique peptides: peptide length, consecutive amino acids that affect trypsin efficiency and/or ragged ends (KK, RK, RR, and KR), chemically active amino acids in the peptide sequence, and peptide locations on the protein tertiary structure.

### The MRM Method Performance

During the analysis, the same sample was injected in the same volume at different time periods. The method of generating proteomics data on biological materials involves several phases. There are currently practical MS quantitation approaches that allow for highly precise quantification in a small or large number of specimens. One of the MS-based proteomics methods, the MRM method (**Table S6**), was used in this investigation.

The performance of the MRM technique is evaluated before studying the behaviors of peptides under various conditions to verify that the method is reproducible. The serum replicates were used to evaluate the run stability of the instrument using Log2 normalized QC serum replicates for each unique peptide in **Figure 3**. The run has decreased and then stabilized over time. **Figure 3A** shows log2 normalized peak areas of peptides in the QC serum sample injected at various times, while **Figure 3B** shows CV values for each target peptide. CV values of each unique peptide do not exceed 2%, and are below 20%, which is accepted in the literature^81–83^. The coefficient of variation of each unique peptide is shown in **Figure 4**. Each target peptide is represented with a different color. When the CV values of the peptides are compared, the unique peptides showing the highest variation are LHTEAQIQEEGTVVELTGR, AAQVTIQSSGTFSSK, and DMYSFLEDMGLK. On the other hand, the target unique peptides with the lowest variation are LPPNVVEESAR and DTVIKPLLVEPEGLEK. Furthermore, the unique peptide LPPNVVEESAR has the highest peak intensity while presenting the least variation. However, peak intensity does not decrease or show same behavior as variation increases.

**Figure 3.**
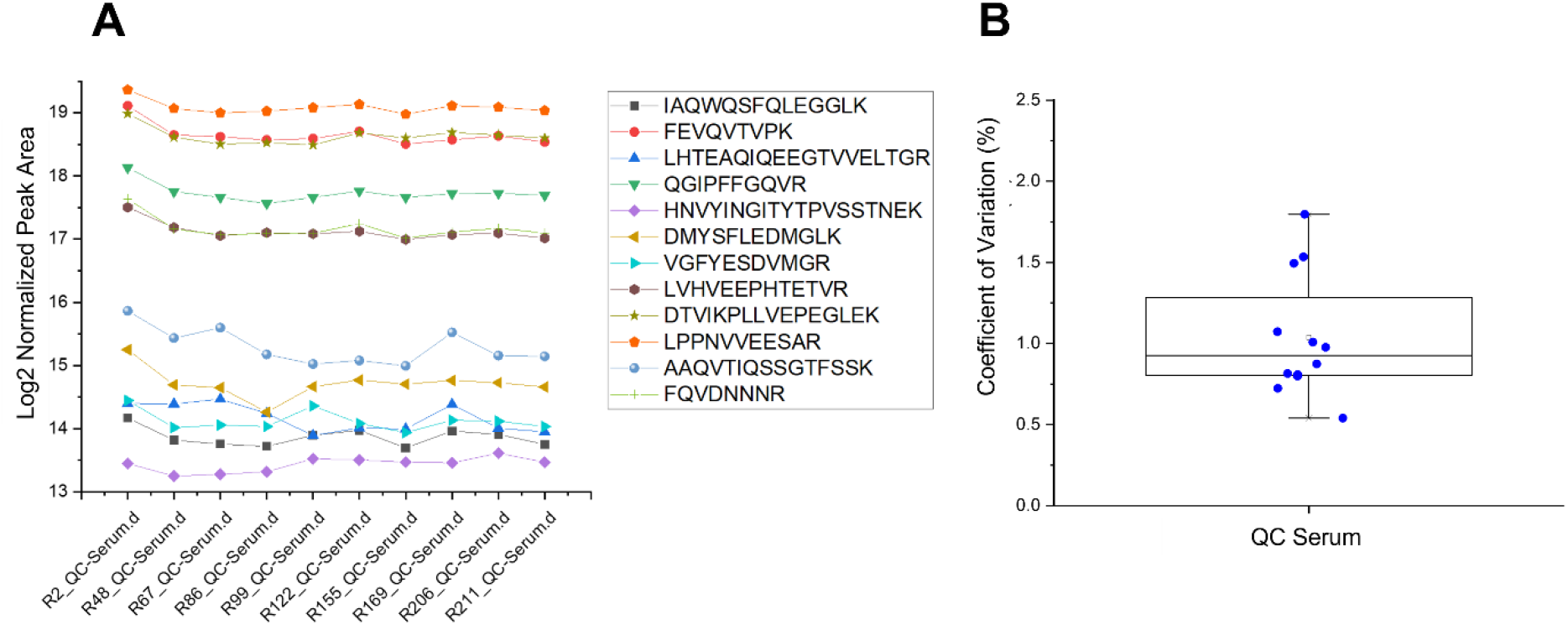
The run stability of QC serum in the QQQ instrument **A**. The normalized peak area of each peptide **B**. CV values of unique peptides in QC serum

**Figure 4.**
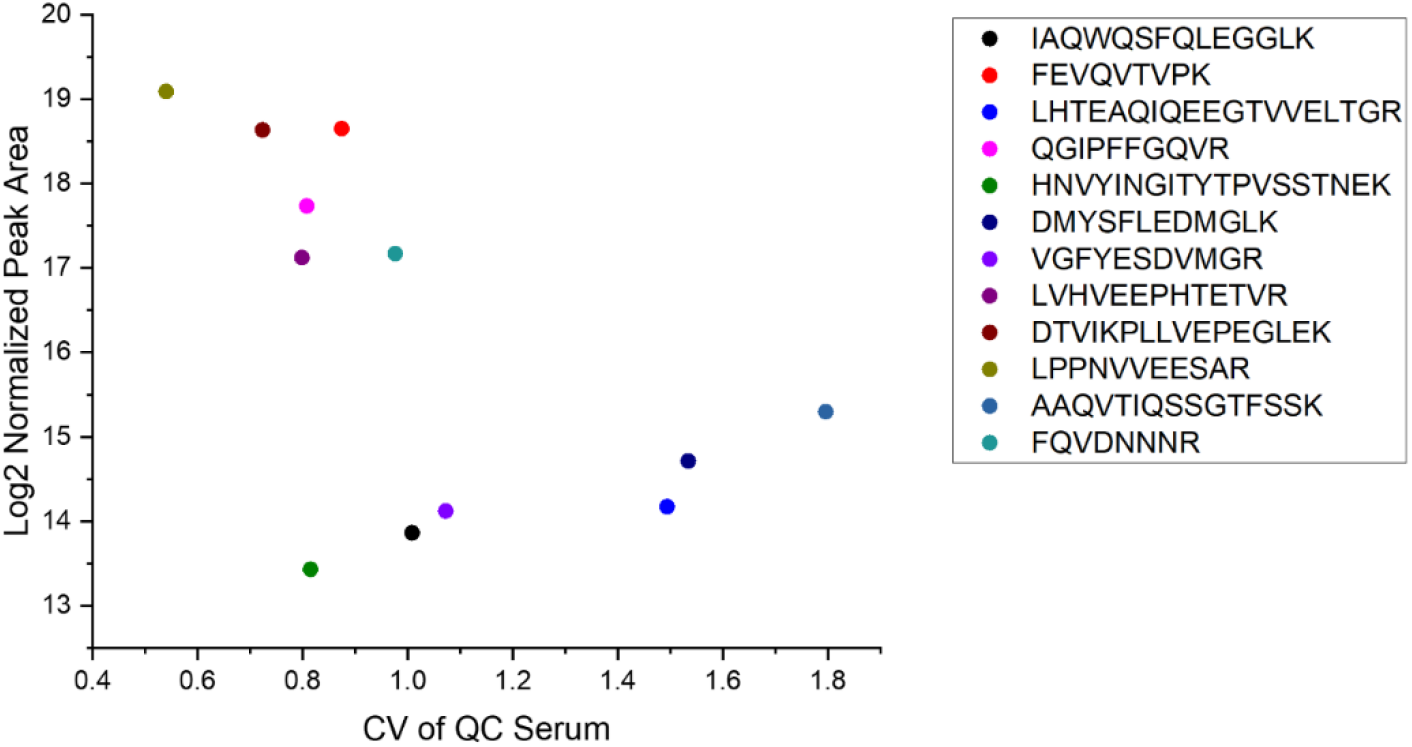
The Log2 normalized peak areas versus CV values of QC serum.

The serum replicates were prepared using the synthetic unique peptide standards. The stability of representative peptide of serum replicates and transitions of its fragments were used to evaluate the method’s performance (**Figure S2**). This process was performed for all peptide standards. **Figure S2** shows the peptide FEVQVTVPK results in Skyline software^84^. The upper window in the figure shows transitions of the unique peptide, and the lower window demonstrates the changes in transition weights of serum replicates and libraries. The stability of serum replicates for a given unique peptide, as well as the consistency of serum replicates, demonstrate that the MRM method is highly reproducible.

### The Digestion Reproducibility

Most recent bottom-up proteomics applications involve quantitative assessment of the absolute and relative protein quantities, affected by several factors such as sample preparation, MS recovery, and data analysis. Protein quantification depends on the efficiency and reproducibility of protein digestion^85^. More specifically, missing or non-specific cleavages in peptide populations have a big impact on protein identification and quantification reliability, as well as repeatability of mass spectrometry analysis^49^. Many non-specific tryptic peptides are considered to be generated during sample preparation. Several factors could account for these nonspecific cleavage events but, the sample preparation has the major impact on the non-specific cleavage. Therefore, the potential to generate reliable digestions that are absent of missing or non-specific cleavages is crucial for protein quantification studies.

Until recently, assessments of digestion products were published at the protein level; thus, studies were complex to the point of being impractical on a proteome-wide scale. However, we focused on protein digestion products at the peptide level. When investigating the digestion reproducibility, log2 normalized serum digest replicates’ results were obtained for both enzymes as shown in **Figure S3**. Trypsin enzyme used replicates, and enzyme mixture results are given in **Figure S3A** and **Figure S3B**. The digestion is reproducible in highly abundant unique peptides for both enzymes, whereas the reproducibility is lower in less abundant unique peptides. The related graphs of coefficient of variation of log2 normalized peak areas of neat human serum samples are given in **Figure S3C** and **Figure S3D**. In both experiments, the CV values are lower than 10%^81–83^. To investigate the coefficient of variation results in detail, CV values of each unique peptide and log2 normalized peak area results were combined. **Figure S4A** illustrates the results of trypsin digestion of neat human serum, while **Figure S4B** shows the results for the use of mixture of enzymes. VGFYESDVMGR, DMYSFLEDMGLK, and FQVDNNNR are the unique peptides with the most variation in the presence of trypsin, as seen in **Figure S4A**. On the other hand, the unique peptides with the largest variation in the enzyme mixture results, as seen in **Figure S4B**, are VGFYESDVMGR, DMYSFLEDMGLK, and HNVYINGITYTPVSSTNEK.

**Figure S4A** shows that when the log2 normalized peak area of unique peptides decreases, the variation increases, and vice versa. The highly abundant peptides have less variation in this case, whereas the less abundant peptides have more significant variation. However, the enzyme mixture result, as shown in **Figure S4B**, does not demonstrate the same behavior observed in **Figure S4A**. It was difficult to specify a relationship between the variation and log2 normalized peak area for unique peptides.

### The Peptide Normalization

Several approaches for peptide normalization that are frequently used in the literature are evaluated. **Figure 5** shows an error bar plot of the twelve unique peptides in six replicates serum digests obtained by using trypsin and trypsin/Lys-C. The error bar plot in **Figure 5A** shows the obtained peak areas of each unique peptide for both enzymes, whereas the second plot shown in **Figure 5B** illustrates the log2 normalization^86^.

**Figure 5.**
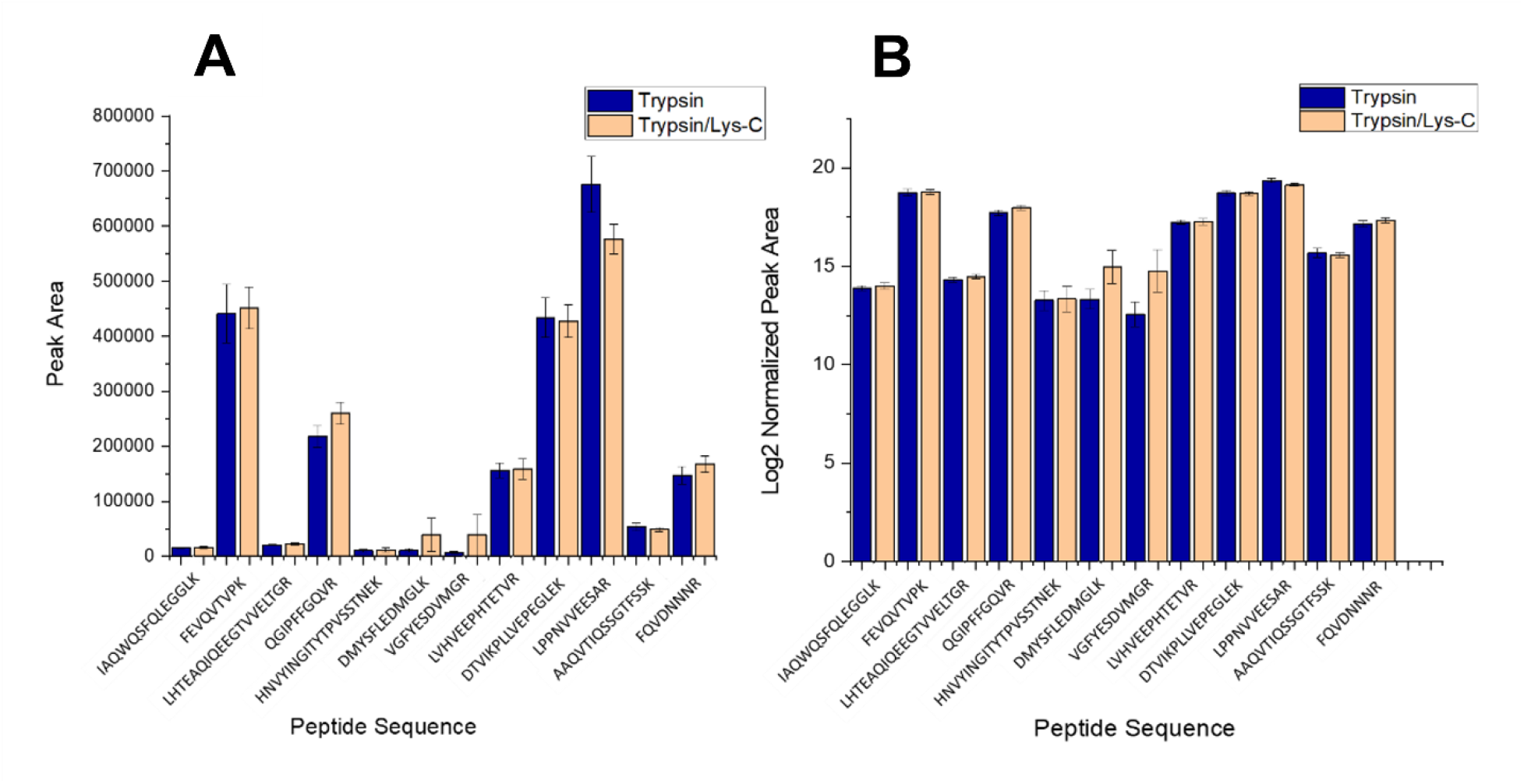
The error column plots of all unique peptides in six serum digest replicates for both enzymes. **A**. The plot of obtained peak areas of unique peptides for both enzymes. **B**. The plot of log2 normalized peak areas of unique peptides for both enzymes.

In order to investigate the root cause of the variation observed in peptide intensities, we examined ionization efficiencies, digestion and matrix effect. The chromatograms of the five unique peptides are shown in **Figure 6**. The equimolar mixture of five synthetic peptides is given in **Figure 6A**. These unique peptides are FEVQVTVPK, QGIPFFGQVR, VGFYESDVMGR, DMYSFLEDMGLK, and LPPNVVEESAR. The variation in peptide intensities are purely related to the ionization efficiencies^87^. The chromatograms of the five unique peptides in the A2MG protein standard are given in **Figure 6B** and **Figure 6C** for trypsin and trypsin/Lys-C, respectively. The relative comparison between target peptides showed variation due to digestion efficiencies. The LPPNVVEESAR unique peptide displayed the greatest intensity decline in the digested samples of A2MG protein in **Figure 6B** and **Figure 6C**, while FEVQVTVPK, VGFYESDVMGR, and DMYSFLEDMGLK unique peptides also displayed a noticeable intensity decrease. The remarkably similar peptide responses in Figures 6B and 6C point to a potential digestion effect. Since DMYSFLEDMGLK peptide cut at the lysine (K) amino acid residue, additional Lys-C is utilized exhibits a change in intensity between the two figures. Furthermore, **Figure 6D** and **Figure 6E** show chromatograms of five unique peptides in neat human serum digested with trypsin and trypsin/Lys-C, respectively. The complex matrix of human serum was digested to produce the figures, and it was shown that QGIPFFGQVR unique peptide responded significantly lower. Besides, the responses of the other four unique peptides analyzed decreased, but not drastically. Thus, the intensity decreases in **Figure 6D** and **Figure 6E** signify ionization and digestion effects as well as matrix effect. The results were compared considering both ionization (shown in **Figure 6A**) and digestion efficiencies (shown in **Figure 6B** and **Figure 6C**). The variation was due to both digestion efficiencies and matrix effect (shown in **Figure 6D** and **Figure 6E**).

**Figure 6.**
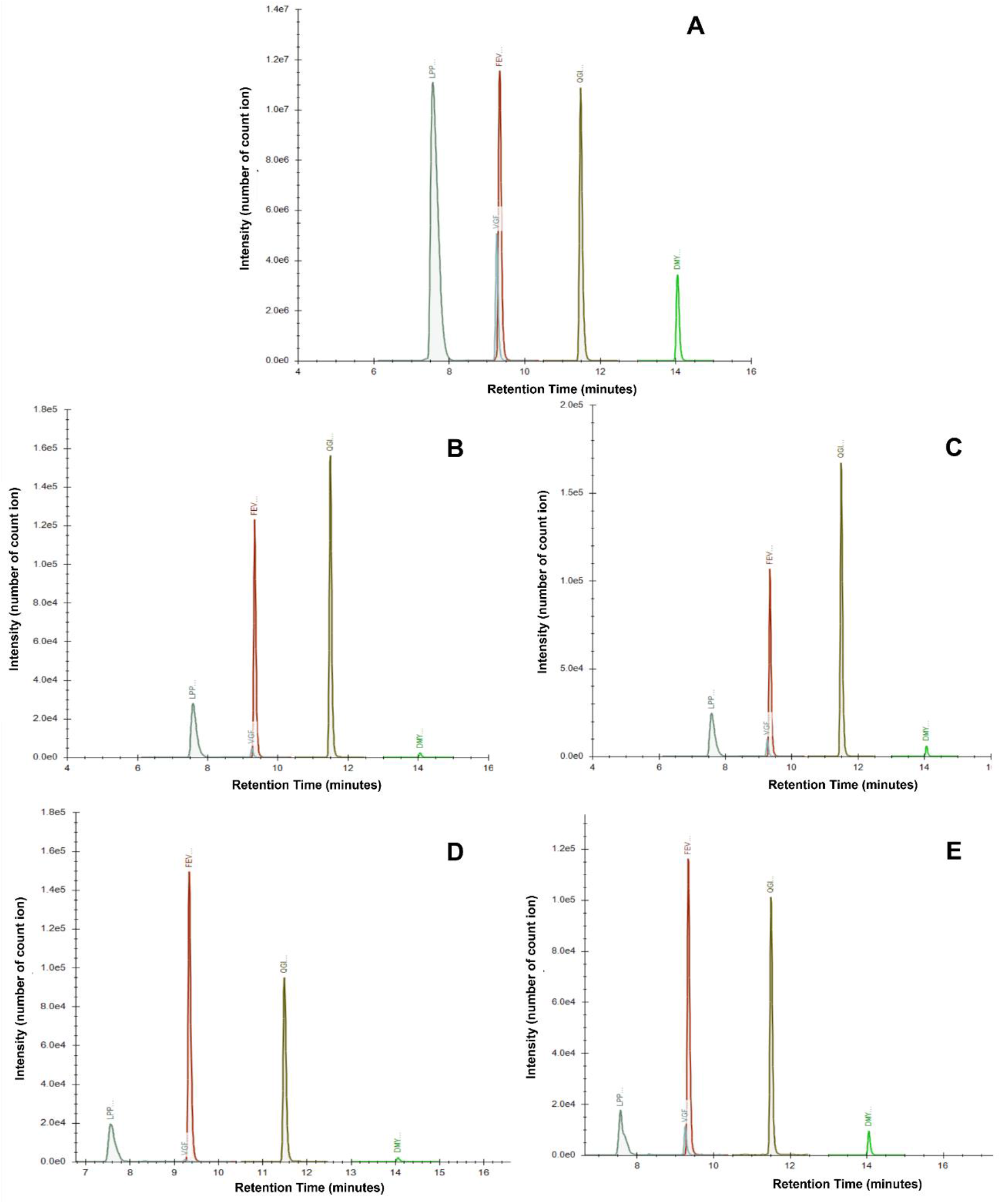
The five unique peptides’ response **A**. in equimolar peptide mixture **B**. in protein standard in the presence of trypsin **C**. in protein standard in the presence of trypsin/Lys-C mixture **D**. in human serum in the presence of trypsin **E**. in human in the presence of trypsin/Lys-C mixture.

The fact that the peptides’ responses change when the enzyme is changed gives support to the claim. Moreover, both digestion and the matrix-effect affected the response change in serum samples. It was observed that the selected unique peptides in the literature were randomly selected either from the highly or the scarcely abundant ones. When assessing digestion reproducibility, the coefficients of variations (CV) of alternative normalizing approaches were also evaluated for both enzyme digestions. Four typical normalization techniques were considered to determine peptide performance, as shown in **Figure 7**.

**Figure 7.**
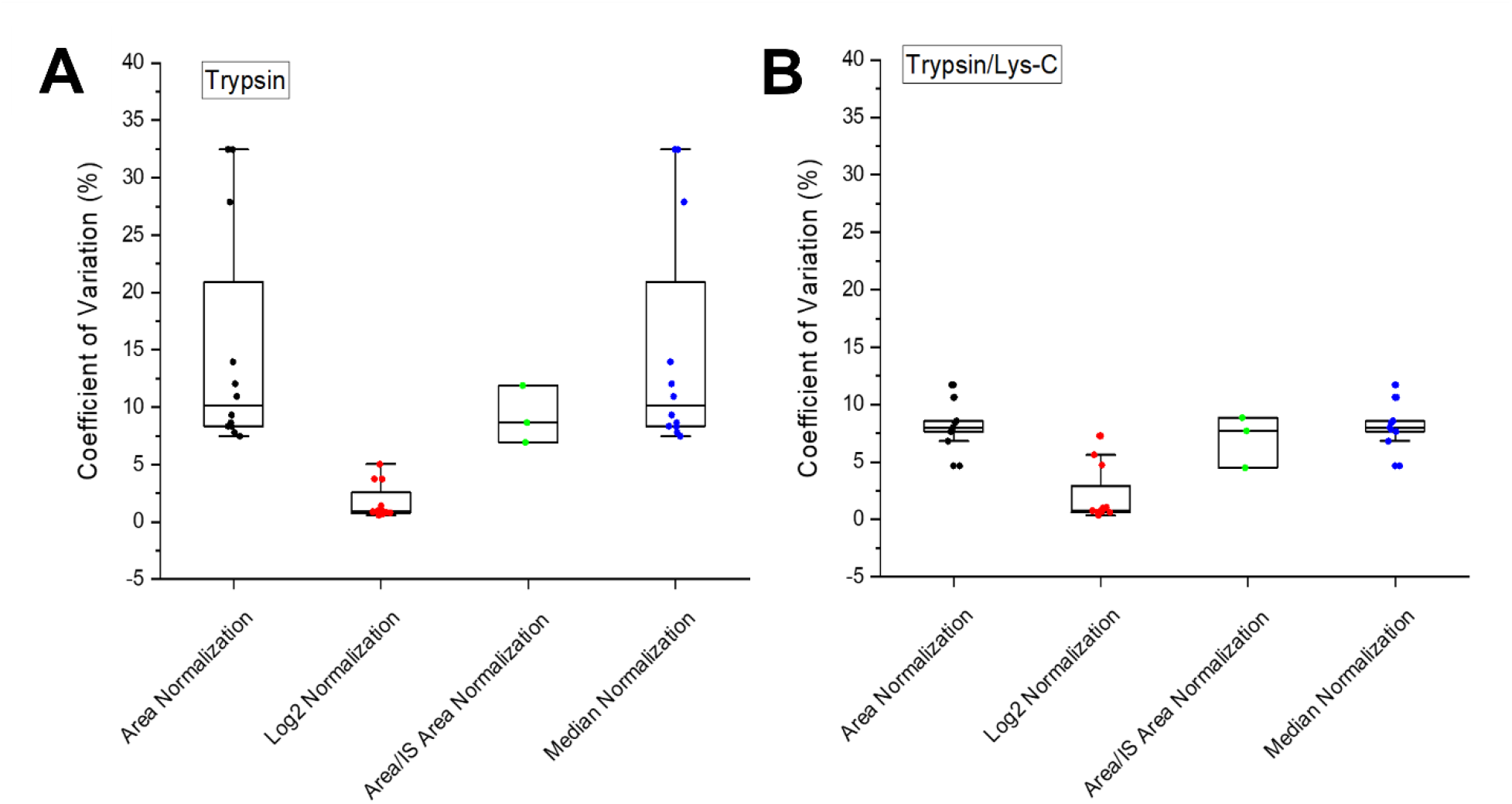
Coefficient of variation (CV) of commonly used normalization approaches in neat human serum for both enzyme digestions. **A**. CV values of different normalization approaches for trypsin digestion **B**. CV values of different normalization approaches for trypsin/Lys-C mixture digestion.

Each data point corresponds to a single peptide in **Figure 7A** and **Figure 7B**. The area, log2, and median normalizations each included twelve unique peptides. Furthermore, the analyte and IS peptide should have a peak area ratio of 1:1. On the other side, the number of protein peptides increases the complexity of optimization. As a result, the literature recommends a ratio of 1:10 to 10:1^32,88,89^ between the sample and the IS peak area for clinical research. However, only three peptides, FEVQVTVPK, DTVIKPLLVEPEGLEK, and LPPNVVEESAR, have the acceptable ratio. In the error bar plots, the whisker range was drawn using a coefficient of 1.5 in respect to the outlier. **Figure 7A** shows that most of the CV values of the peptides in the trypsin digestion graph are under 10%, and two peptides have more than 20% variance. **Figure 7B** shows the results of the trypsin/Lys-C digestion, which show that all the CV values are less than 20% variation. These CV values indicate that each target peptide has less variance. When the analytical variation is less, the biological variation is observed more clearly. Although some sources suggest that CV values should be less than 10%^90–92^, the literature indicates that the threshold is 20%^81–83^.

### The Effect of Concentration

The concentration effect of A2MG protein standard on unique peptides was visualized using heatmaps with a dendrogram graph. To investigate the impact of concentration, A2MG protein standard samples were compared to one another. For the comparison to be valid, Pearson correlated plots were drawn in the same correlation scales (between -1 to +1), shown in **Figure S5**. When evaluating the concentration effect, A2MG protein samples prepared in eight concentration levels were used. The graphs in **Figure S5** are divided into two categories based on the concentration of protein standard samples: high concentration and low concentration. The low concentration A2MG protein standard samples shown in **Figure S5A** and **Figure S5B** have a concentration range of 0.0071 – 0.0357 μg/μl, whereas the high concentration protein standards demonstrated in **Figure S5C** and **Figure S5D** have a concentration range of 0.0536 – 0.1071 μg/μl. When the cluster analysis is evaluated, the dendrograms of the four plots in **Figure S5** are divided into two major subgroups. Any association between the location and abundance of the peptides included in these clusters was not found when the subgroups were examined. When we compare high (**Figure S5B**) and low concentration (**Figure S5D**) of A2MG protein standard samples digested by trypsin and trypsin/Lys-C mixture, the linear correlation is found to be more significant at high concentration. The opposite linear correlation appears to be stronger in protein standard samples with low concentration (**Figure S5A** and **Figure S5B**) than in protein standard samples with high concentration (**Figure S5C** and **Figure S5D**). This weaker signal may have resulted in a different correlation. Apart from the unique peptide HNVYINGITYTPVSSTNEK, it is observed that eleven unique peptides behave differently in the presence of trypsin at low protein concentrations. Also, the unique peptide LHTEAQIQEEGTVVELTGR in the presence of a trypsin/Lys-C mixture at low protein concentration levels is not correlated with other unique peptides. These differences in correlation may be due to insufficient information gathered due to poor MS signal response of protein standard samples at low concentrations. When the correlation of protein standard samples digested with trypsin is compared in **Figure S5A** and **Figure S5C**, it is seen that samples with higher concentrations have a higher correlation. However, the unique peptide DMYSFLEDMGLK and VGFYESDVMGR show opposite linear correlations. Similarly, in **Figure S5B** and **Figure S5D**, the unique peptides in the presence of trypsin/Lys-C mixture are correlated better in high concentration protein standard samples. Once again, trypsin/Lys-C mixture provides a successful cleavage in the inner regions of the protein and better digestion than trypsin. On the other hand, it appears to have a weaker opposite linear correlation with the unique peptides, DMYSFLEDMGLK, AAQVTIQSSGTFSSK, and VGFYESDVMGR. The strong correlation could be attributed to the fact that these peptides are located very close to glycosylation sites in the protein structure. On the other hand, the unique peptides’ active amino acid residues such as methionine (M), glutamine (Q), and histidine (H) could account for the weak correlation. Thus, the unique peptide LHTEAQIQEEGTVVELTGR would have an opposite correlation with the unique peptide VGFYESDVMGR but a weaker opposite correlation with the unique peptides LPPNVVEESAR, LVHVEEPHTETVR, QGIPFFGQVR, DTVIKPLLVEPEGLEK, HNVYINGITYTPVSSTNEK, and FQVDNNNR in **Figure S5B**. This relationship could have occurred between peptides whose digestion was incomplete. Since the peptide LHTEAQIQEEGTVVELTGR is one of the longest peptides, an increment in peptide length might be a reason for incomplete digestion. Furthermore, this unique peptide has consecutive amino acid residues such as EK or RK, that might be reducing the complete digestion. Based on these results, it can be deduced that digestion of this unique peptide is incomplete and that it has a relation with the peptides with which it is oppositely correlated through this factor. In addition, the highest correlation was observed in high concentration levels of protein standard samples digested with trypsin/Lys-C mixture (**Figure S5D**). While all peptides correlate with each other in **Figure S5D**, the unique peptides in high concentration levels of protein standard digested with trypsin shows an opposite correlation in **Figure S5C**. These unique peptides showing opposite correlation are DMYSFLEDMGLK and VGFYESDVMGR in **Figure S5C**, and **Figure S6** shows a zoomed-in version of the heatmap with a dendrogram graph for these peptides. In the coefficient of variation (**Figure S6A**), these unique peptides likewise had the highest CV values, which are marked in **Figure S6B**. These are the peptides with the lowest pI values (3.54 and 3.93, respectively) and are found within the protein core region. Furthermore, they have nearly the same peptide length (12 and 11, respectively). The findings lead to the following conclusion: a low pI value may be a factor that negatively impacts trypsin’s function, resulting in poor digestion in the protein’s inner regions or negatively affects the ionization of unique peptides.

### The Effect of Enzymes on Digestion Process

The effect of different enzymes on the digestion process was investigated by comparing digestion of A2MG protein standard and human serum samples by trypsin and trypsin/Lys-C mixture. **Figure 8** depicts all heatmaps with dendrogram using Pearson’s correlation on the same scale. According to the hierarchical cluster analysis, the included protein standard and serum samples were separated into two classes. High concentration levels of the A2MG protein standard were utilized to create this heatmap graph, as well as three concentration levels generated by spiking A2MG protein into the human serum and neat human serum. **Figure 8A** and **Figure 8B** represent the correlation of peptides in protein standard samples, while **Figure 8C** and **Figure 8D** represent the correlation in standard serum samples. Thus, the comparison in the pure protein standard and serum sample with complex matrix has been investigated. Trypsin and trypsin/Lys-C mixture has been used to compare digestion efficiency with missed cleavage. The highest correlation was observed in protein standard samples. The peptides VGFYESDVMGR and DMYSFLEDMGLK have a complete opposite correlation with other peptides as shown in **Figure 8A**. They are located at similar sites in the 3D structure. The peptide VGFYESDVMGR is in the bait region (described as core region), whereas the peptide DMYSFLEDMGLK is located at the inner region of the protein. This result exhibits that these two peptides have very similar properties and are different from other unique peptides. When **Figure 8A** and **Figure 8B** are compared, the dissimilarity coefficient decreases for trypsin/Lys-C mixture digestion compared to trypsin digestion. The main reason for this differentiation could be the location of these longer peptides.

**Figure 8.**
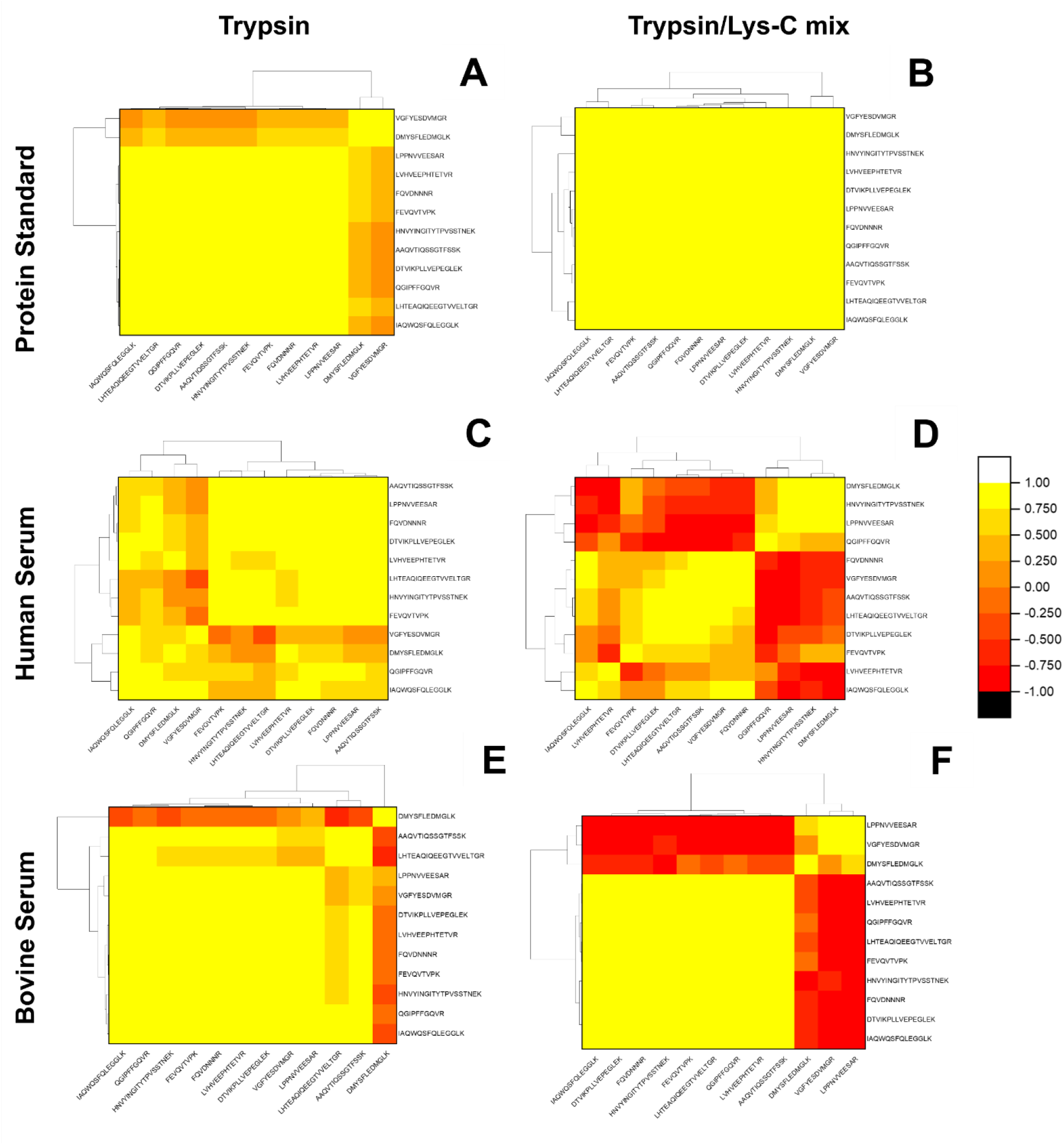
A heatmap with a dendrogram plot illustrating the association of A2MG unique peptides in human and bovine serum-protein spiked samples in the presence of various enzymes. **A**. Peptide correlation in the presence of trypsin in protein standard. **B**. The correlation in the presence of trypsin/Lys-C mixture in protein standard. **C**. The correlation in the presence of trypsin in bovine serum **D**. The correlation in the existence of trypsin/Lys-C mixture in bovine serum **E**. The correlation in the presence of trypsin in human serum. **F**. The correlation in the presence of trypsin/Lys-C mixture in human serum.

Trypsin/Lys-C mixture performs a better cleavage because enzymes can reach the inner regions considerably better than only trypsin and prevent missed cleavages. Besides, when all unique peptides in **Figure 8A** and **Figure 8B** are compared, the decrease in linear correlation validates our hypothesis. On the other hand, serum digest samples’ peptide correlations are not as simple as protein standard samples because of higher complexity, including other proteins and different forms of proteins. When the increase in environment’s heterogeneity, VGFYESDVMGR and DMYSFLEDMGLK peptides are still highly opposite correlated as shown in **Figure 8C**, though correlation was not as substantial as the correlation displayed in **Figure 8A**.

The unique peptides HNVYINGITYTPVSSTNEK, LHTEAQIQEEGTVVELTGR, and LVHVEEPHTETVR, are correlated with each other as shown in **Figure 8C**. The reason for the linear correlation may be the presence of histidine (H) residues in these sequences. The chemical modifications can cause a change in peptide forms^93^. A modified form of these peptides may have occurred, which may have also affected the peptides’ correlation.

### The Peptide Behavior in Different Matrices

To investigate the behavior of the A2MG protein’s unique peptides in various matrices, the bovine serum matrix was used as an alternative matrix to the human serum matrix. The reason is that the bovine serum is easily available, has a complex environment as human serum, and does not contain all of the unique peptides of A2MG protein found in human sera. **Table S7** below lists the unique peptides in the human A2MG protein as well as the same peptides in the bovine A2MG protein. In this part of the analysis, bovine serum and human serum were spiked with A2MG protein standard and digested with trypsin enzyme and a trypsin/Lys-C mixture. The serum samples are compared with protein standards. A2MG concentrations in spiked A2MG protein into human serum range from 0.0455 to 0.1000 μg/μl, in the A2MG protein standard from 0.0536 to 0.1071 μg/μl, and spiked A2MG protein into bovine serum from 0 to 0.0121 μg/μl. In addition, A2MG concentration was considered nonexistent in bovine serum since it does not contain all of the human specific A2MG unique peptides.

In bovine serum samples, there is an opposite correlation. Furthermore, the opposite correlation in unique peptides in trypsin digested samples was lower than in the trypsin/Lys-C digested samples (**Figure 8C** and **Figure 8D**, respectively). Likewise, the opposite correlation was stronger in unique peptides in presence of the trypsin/Lys-C digested samples than in the trypsin digested human serum samples in **Figure 8E** and **Figure 8F**, respectively. When all of the plots in **Figure 8** are examined, the protein standard plots exhibit the strongest linear peptide correlation compared to the serum plots. When the graphs are evaluated over the digestion process, it can be seen that the enzyme mixture results are the strongest positive or negative peptide correlation than the trypsin enzyme.

When human and bovine serum samples are compared, human serum samples are correlated better than bovine serum samples because they already contain these twelve unique peptides of the A2MG protein. Thus, the peptides’ concentration in human serum is higher than in bovine serum, revealing a better linear correlation. **Figure 8C** shows that the unique peptide DMYSFLEDMGLK has a high opposite correlation with other A2MG unique peptides. The unique peptide DMYSFLEDMGLK with one of the lowest pI values is found inside the A2MG protein. Although the linear correlation in human serum samples is higher than in bovine serum samples in the presence of trypsin enzyme, the unique peptides DMYSFLEDMGLK and VGFYESDVMGR differ from other unique peptides found in trypsin digested human serum samples, as shown in **Figure 8E**. However, trypsin digested human serum samples have the highest linear correlation of these four serum plots in **Figure 8**. The highly opposite correlations of three unique peptides, DMYSFLEDMGLK, LPPNVVEESAR, and VGFYESDVMGR, are demonstrated in **Figure 8D**. These three unique peptides’ lengths are 12, 11, and 11, respectively. The active methionine (M) amino acid residue is found in the peptides DMYSFLEDMGLK and VGFYESDVMGR. This residue may be a distinguishing factor since only these two unique peptides have this active amino acid residue among the twelve unique peptides. In addition, these three unique peptides have extremely low pI values among the twelve unique peptides of A2MG, which are 3.54, 3.93, and 4.15, correspondingly. As a result, it may be one of the major factors driving this correlation. Furthermore, since the unique peptides DMYSFLEDMGLK and VGFYESDVMGR are found in the inner region of the protein, even the unique peptide VGFYESDVMGR is located in the bait region. Thus, incomplete digestion may be one of the factors affecting the correlation. The variation in peptide level might be affected by various parameters such as the presence of chemically active amino acids in the sequence, the ragged peptide ends that impact activity of trypsin, peptide length, pI values, the locations in the protein structure and potential modifications on peptide sequence.

## CONCLUSION

There is a growing interest in high-throughput protein analysis using both untargeted and targeted proteomics methods. Furthermore, there are huge efforts developing quantitative proteomics assays for clinical applications. However, these analyses target often limited number of unique peptides for each protein. The analytical performance of the methods is generally measured based on the variation of a single peptide. Although this approach is applicable for relative comparison between groups, it may not reflect the biological changes occurring in protein level. The question rises whether multiple unique peptides representing same protein are correlated to each other or not. Until now, the studies using two or more peptides for the same protein yielding different results were linked to ionization efficiencies. They are often classified as outliers and eliminated from the studies. To this end, we used a systematic approach to show dynamic protein-peptide correlation by monitoring 12 unique peptides at varying concentrations of the protein. Results suggest that the change in A2MG protein concentration at protein level is not directly reflected in the unique peptide level as responses were different. Consequently, the selection of a unique peptide for any clinical application is critical as one could get different results depending on the choice. While any unique peptide could be sufficient enough to identify/discover the protein of interest in the sample, as it is shown peptide selection is a crucial step for protein quantification. Replicate measurements showed a reproducible protein digestion and MS analysis for majority of the peptides. This is a misleading control measure and as in our study, they were not correlated to each other. This leads us reported results indicate peptide-based markers rather than protein. In order to assess the dynamic protein-peptide relationship considering ionization, digestion, matrix and concentration perspectives, this study is limited to one representative protein and single digestion procedure. Future studies need to investigate more proteins and processing conditions to understand underlying processing and biological factors.

## Table of Contents (TOC) Graphic / Abstract Graphic

(8.25 cm × 4.28 cm)

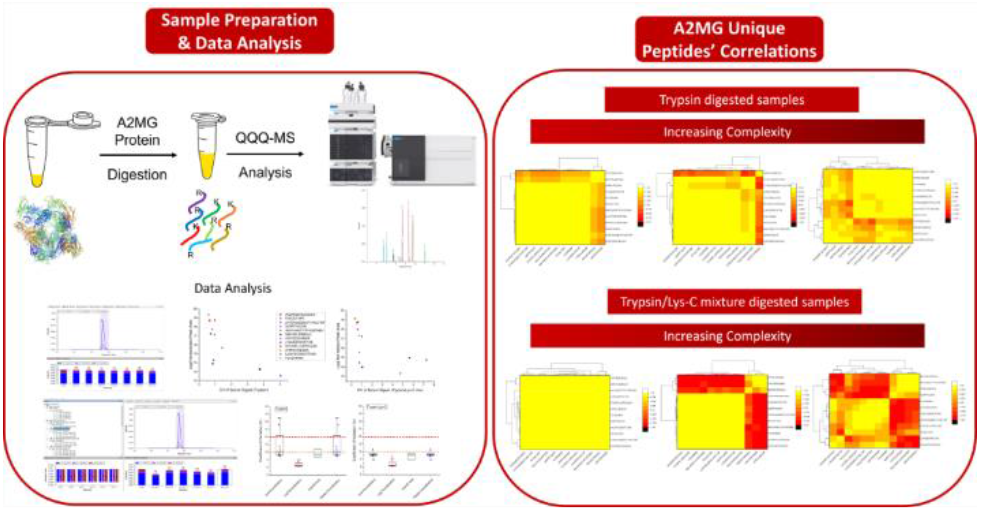

## ASSOCIATED CONTENT

The Supporting Information is available.

**Figure S1**. Positions of lysine and arginine in the A2MG protein. The purple color code is used for lysine residues and the yellow color code for the arginine residues. **Table S1**. The list and information of unique tryptic peptides in A2MG protein. **Table S2**. The criteria for identifying unique representative peptides. **Table S3**. The color codes of unique peptides in **Figure 1A**. Table S4. The list of unique peptides located near to glycosylation. **Table S5**. The number of publications of A2MG unique peptides in literature. **Table S6**. The list of selected MRM transitions of unique peptides of A2MG protein. **Figure S2**. The replicate stability of unique peptide FEVQVTVPK of QC serum in Skyline software. **Figure S3**. The replicates of standard neat human serum samples for both enzymes. A. In the presence of trypsin, each peptide’s normalized response is reported in replicates of neat human serum. B. In the presence of trypsin/Lys-C mixture enzyme, each peptide’s normalized response is reported in replicates of neat human serum. C. CV values of peptides digested with trypsin D. CV values of peptides digested with trypsin/Lys-C mixture. **Figure S4**. The Log2 normalized peak areas versus CV values of neat human serum samples. A. in trypsin B. in trypsin/Lys-C mixture. **Figure S5**. A heat map with a dendrogram plot demonstrating the association of A2MG unique peptides in protein standard samples presents different enzymes. A. Correlation in low concentration protein standard samples in the presence of trypsin. B. Correlation in low-concentration protein standard samples with trypsin/Lys-C mixture. C. The correlation in high concentration protein standards presence of trypsin. D. Correlation in high-concentration protein standard samples with trypsin/Lys-C mixture. **Figure S6**. The oppositely correlated peptides in Figure S5C with the highlighted CV values. A. The peptide correlation due to trypsin digestion in Figure S5C. The oppositely correlated peptides are highlighted. B. The CV values of these peptides are highlighted. **Table S7**. List of A2MG protein’s unique peptides shared to both matrices. *The peptide has not been included due to low ionization during MS analysis **The synthetic unique peptides were purchased.

## AUTHOR INFORMATION

### Author Contributions

The manuscript was written through contributions of all authors. All authors have given approval to the final version of the manuscript.

### Notes

The authors declare no competing financial interest.

## ACKNOWLEDGMENT

We are extremely grateful to all the participants who took part in this study. This work was supported by grants from the Middle East Technical University Scientific Research Projects Coordination Unit (METU BAP AGEP-103-2019-10105).

## ABBREVIATIONS

A2MG: alpha-2-macroglobulin
ABC: ammonium bicarbonate
ACN: acetonitrile
CPTAC: Clinical Proteomic Tumor Analysis Consortium
CV: coefficient of variation
DTT: dithiothreitol
ESI: electrospray ionization
IAA: iodoacetamide
IS: internal standard
LC: liquid-chromatography
Lys-C: lysine-C
MS: mass spectrometry
MRM: multiple reaction monitoring
m/z: mass-to-charge ratio
PDB: Protein Database Bank
Q: quadrupole
QC: quality control
QQQ: triple quadrupole.

