## Supplementary Data for "A single protein to multiple peptides: Investigation of protein-peptide relationship using targeted alpha-2-macroglobulin analysis"

***Corresponding Author:**

Assist. Prof. Dr. Sureyya Ozcan

orcid.org/0000-0002-9371-3696

**TABLE OF CONTENTS**

**Figure S1.** Positions of lysine and arginine in the A2MG protein. The purple color code is used for lysine residues and the yellow color code for the arginine residues.................................................................S2

**Table S1.** The list and information of unique tryptic peptides in A2MG protein…………………………S3

**Table S2.** The criteria for identifying unique representative peptides…………………………………….S4

**Table S3.** The color codes of unique peptides in **Figure 1A**……………………………………………..S5

**Table S4.** The list of unique peptides located near to glycosylation……………………………………..S5

**Table S5.** The number of publications of A2MG unique peptides in literature…………………………..S6

**Table S6.** The list of selected MRM transitions of unique peptides of A2MG protein……………………S7

**Table S7.** List of A2MG protein’s unique peptides shared to both matrices. *The peptide has not been included due to low ionization during MS analysis **The synthetic unique peptides were purchased...........................................................................................................................................S11

**References**…...………………………………………………………………………………………………S12

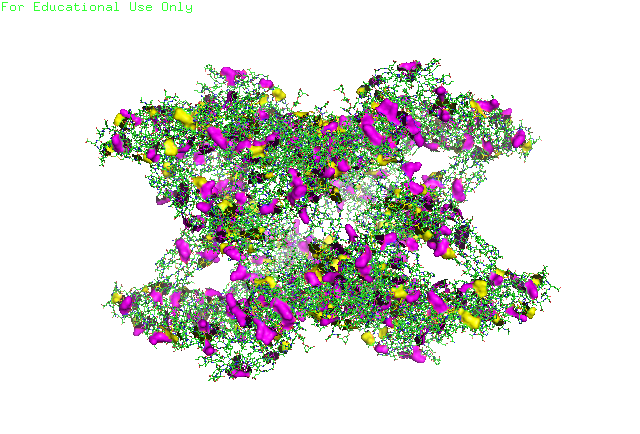

**Figure S1.** Positions of lysine and arginine in the A2MG protein. The purple color code is used for lysine residues and the yellow color code for the arginine residues.

**Table S1.** The list and information of unique tryptic peptides in A2MG protein

| **Position of the cleavage** | **Position of Peptide**  **Sequence** | **Peptide Sequence** | **Peptide Length (Number of**  **amino acids)** |
| --- | --- | --- | --- |
| 188 | 175-188 | IAQWQSFQLEGGLK | 14 |
| 237 | 229-237 | FEVQVTVPK | 9 |
| 370 | 361-370 | QGIPFFGQVR | 10 |
| 338 | 320-338 | LHTEAQIQEEGTVVELTGR | 19 |
| 664 | 646-664 | HNVYINGITYTPVSSTNEK | 19 |
| 676 | 665-676 | DMYSFLEDMGLK | 12 |
| 715 | 705-715 | VGFYESDVMGR | 11 |
| 732 | 720-732 | LVHVEEPHTETVR | 13 |
| 863 | 854-863 | QTVSWAVTPK | 10 |
| 912 | 897-912 | DTVIKPLLVEPEGLEK | 16 |
| 945 | 935-945 | LPPNVVEESAR | 11 |
| 1122 | 1093-1122 | GGVEDEVTLSAYITIALLEIPLTVTHPVVR | 30 |
| 1162 | 1148-1162 | ALLAYAFALAGNQDK | 15 |
| 1289 | 1275-1289 | AAQVTIQSSGTFSSK | 15 |
| 1297 | 1290-1297 | FQVDNNNR | 8 |

**Table S2.** The criteria for identifying unique representative peptides

| **Peptide Sequence** | **Peptide Length** | **Peptide Position** | **Sequential AA**  **affecting trypsin efficiency** | **Chemically Active**  **Amino Acid Residues** | | | | | |
| --- | --- | --- | --- | --- | --- | --- | --- | --- | --- |
|  |  |  |  | **Methionine (M)** | **Cysteine (C)** | **Glutamine (Q)** | **Tryptophan (W)** | **Histidine (H)** | **Asparagine (N)** |
| IAQWQSFQLEGGLK | 14 | 175-188 |  | 0 | 0 | 3 | 1 | 0 | 0 |
| FEVQVTVPK | 9 | 229-237 | K/E | 0 | 0 | 1 | 0 | 0 | 0 |
| LHTEAQIQEEGTVVELTGR | 19 | 320-338 | EK, R/K | 0 | 0 | 2 | 0 | 1 | 0 |
| QGIPFFGQVR | 10 | 361-370 |  | 0 | 0 | 2 | 0 | 0 | 0 |
| HNVYINGITYTPVSSTNEK | 19 | 646-664 |  | 0 | 0 | 0 | 0 | 1 | 3 |
| DMYSFLEDMGLK | 12 | 665-676 |  | 2 | 0 | 0 | 0 | 0 | 0 |
| VGFYESDVMGR | 11 | 705-715 |  | 1 | 0 | 0 | 0 | 0 | 0 |
| LVHVEEPHTETVR | 13 | 720-732 | R/K | 0 | 0 | 0 | 0 | 2 | 0 |
| DTVIKPLLVEPEGLEK | 16 | 897-912 | RK | 0 | 0 | 0 | 0 | 0 | 0 |
| LPPNVVEESAR | 11 | 935-945 |  | 0 | 0 | 0 | 0 | 0 | 1 |
| ALLAYAFALAGNQDK | 15 | 1148-1162 | K/RK | 0 | 0 | 1 | 0 | 0 | 1 |
| AAQVTIQSSGTFSSK | 15 | 1275-1289 |  | 0 | 0 | 2 | 0 | 0 | 0 |
| FQVDNNNR | 8 | 1290-1297 |  | 0 | 0 | 1 | 0 | 0 | 3 |

**Table S3.** The color codes of unique peptides in **Figure 1A**

| **Sequence**  **Position** | **Peptide Sequence** | **Color Name** | **Color Code** |
| --- | --- | --- | --- |
| 175-188 | IAQWQSFQLEGGLK | Red |  |
| 229-237 | FEVQVTVPK | Yellow |  |
| 320-338 | LHTEAQIQEEGTVVELTGR | Lilac |  |
| 361-370 | QGIPFFGQVR | Pale Blue |  |
| 646-664 | HNVYINGITYTPVSSTNEK | Cyan |  |
| 665-676 | DMYSFLEDMGLK | Pink |  |
| 705-715 | VGFYESDVMGR | - | - |
| 720-732 | LVHVEEPHTETVR | - | - |
| 897-912 | DTVIKPLLVEPEGLEK | Blue |  |
| 935-945 | LPPNVVEESAR | Olive |  |
| 1148-1162 | ALLAYAFALAGNQDK | Violet |  |
| 1275-1289 | AAQVTIQSSGTFSSK | Sea Blue |  |
| 1290-1297 | FQVDNNNR | Salmon |  |

**Table S4.** The list of unique peptides located near to glycosylation

| **Peptide Sequence** | **Peptide Length** | **Peptide Position** | **PTM** |
| --- | --- | --- | --- |
| FEVQVTVPK | 9 | 229-237 | Next to Glycosylation at 247 |
| QGIPFFGQVR | 10 | 361-370 | Next to Glycosylation at 410 |
| DTVIKPLLVEPEGLEK | 16 | 897-912 | Next to Glycosylation at 869 |

**Table S5.** The number of publications of A2MG unique peptides in literature

| **The Number of Publications** | **Peptide Sequence** |
| --- | --- |
| 13 ^1–13^ | IAQWQSFQLEGGLK |
| 18 ^1–11,14–20^ | FEVQVTVPK |
| 13 ^1,3–6,9–13,15,21,22^ | LHTEAQIQEEGTVVELTGR |
| 18 ^2–7,9–15,17,18,23–25^ | QGIPFFGQVR |
| 10 ^1–4,6,11–13,26,27^ | HNVYINGITYTPVSSTNEK |
| 11 ^1–4,6,10–13,18,23^ | DMYSFLEDMGLK |
| 13 ^1–6,9–14,18^ | VGFYESDVMGR |
| 14 ^1–6,11–15,18,24,25^ | LVHVEEPHTETVR |
| 15 ^1–7,9–11,13–15,18,23^ | DTVIKPLLVEPEGLEK |
| 18 ^1–9,11–14,17–19,23,28^ | LPPNVVEESAR |
| 11 ^1,3,5,6,8,9,11,13,15,17,19^ | AAQVTIQSSGTFSSK |
| 9 ^3,5–9,11,12,14^ | FQVDNNNR |

**Table S6.** The list of selected MRM transitions of unique peptides of A2MG protein

| **Protein UniProt ID** | **Peptide Sequence** | **Retention Time (min)** | **Predominant Charge State** | **Precursor Ion (m/z)** | **Transition ion (y,b)** | **Transitions (m/z)** | **Collision Energy (V)** |
| --- | --- | --- | --- | --- | --- | --- | --- |
| Alpha-2-macroglobulin P01023 (A2MG_HUMAN) | IAQWQSFQLEGGLK | 10.9 | +3 | 535.62 | y7 y6 b10 | 744.42+ 616.37+ 616.31+ | 14.5 |
|  | FEVQVTVPK | 9.4 | +2 | 523.71 | y7 y6 y5 b6 | 770.48+ 671.49+ 543.35+ 704.36+ | 14.1 |
|  | LHTEAQIQEEGTVVELTGR | 8.8 | +3 | 704.03 | y9 y7 b7 b15 | 931.52+ 773.45+ 793.42+ 832.91++ | 20.5 |
|  | QGIPFFGQVR | 11.5 | +2 | 574.81 | y7 y6 y5 y7 | 850.46+ 753.40+ 606.34+ 425.73++ | 15.9 |
|  | HNVYINGITYTPVSSTNEK | 8.6 | +3 | 713.02 | y9 y8 y8 b11 | 962.48+ 861.43+ 431.22++ 638.82++ | 20.9 |
|  | DMYSFLEDMGLK | 6.1 | +2 | 724.83 | y9 y8 y7 | 1039.52+ 952.48+ 805.41+ | 21.3 |
|  | VGFYESDVMGR | 14 | +2 | 630.29 | y10 y8 y7 y6 | 1160.50+ 956.41+ 793.35+ 664.31+ | 17.9 |
|  | LVHVEEPHTETVR | 9.3 | +3 | 515.94 | y11 y10 y7 b6 | 667.33++ 598.80++ 420.22++ 707.37+ | 13.8 |
|  | DTVIKPLLVEPEGLEK | 10.7 | +3 | 594.01 | y6 y6 y14 y13 | 672.36+ 336.68++ 782.47++ 732.94++ | 16.6 |
|  | LPPNVVEESAR | 7.6 | +2 | 605.83 | y6 y10 y9 y9 | 690.34+ 549.28++ 1000.51+ 500.76++ | 17 |
|  | AAQVTIQSSGTFSSK | 7.5 | +2 | 756.39 | y11 y10 y9 y8 | 1142.57+ 1041.52+ 928.44+ 800.38+ | 22.4 |
|  | FQVDNNNR | 5.5 | +2 | 503.74 | y7 y6 y5 y4 | 859.40+ 731.34+ 632.28+ 517.25+ | 13.3 |

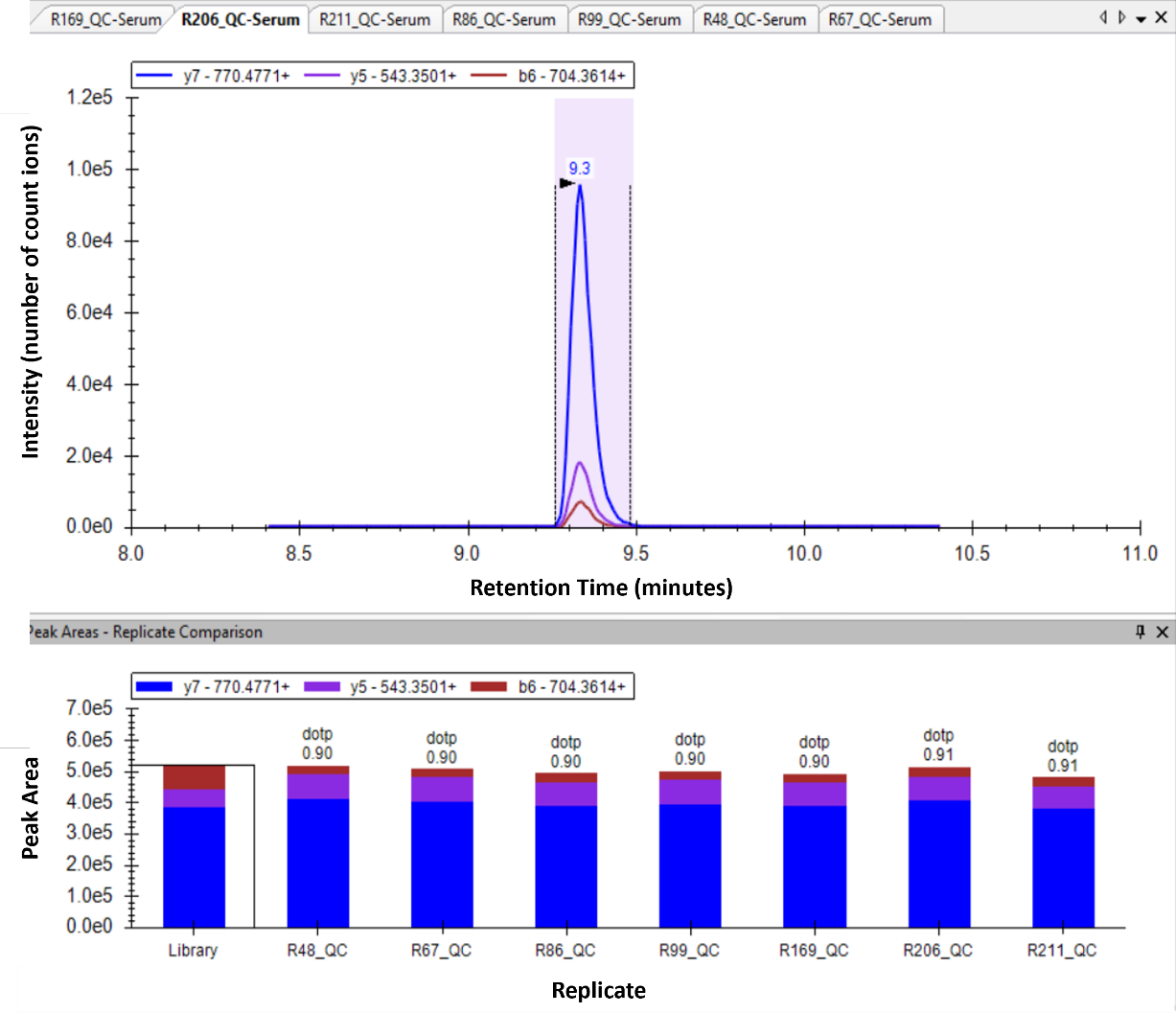

**Figure S2.** The replicate stability of unique peptide FEVQVTVPK of QC serum in Skyline software.

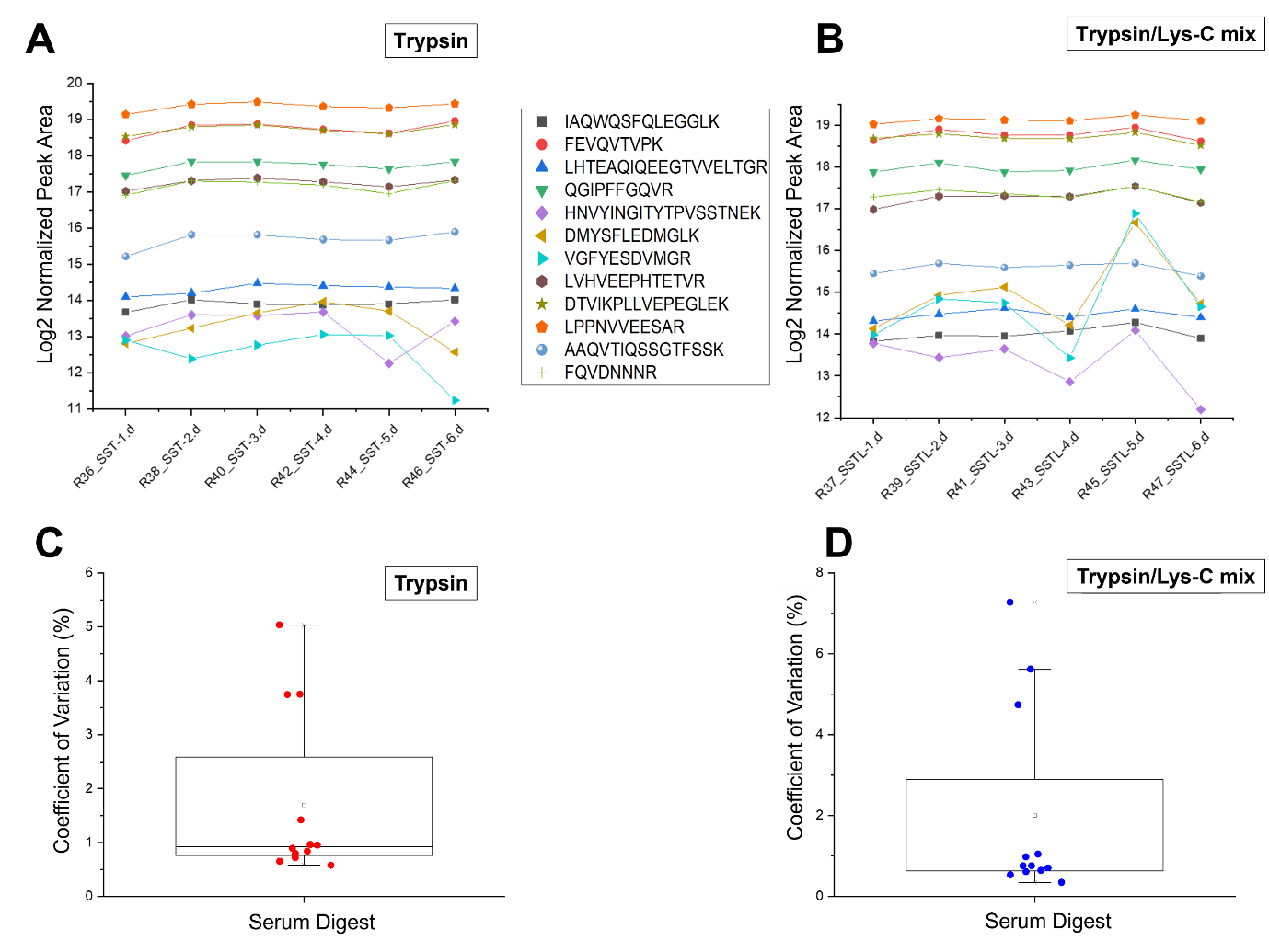

**Figure S3.**The replicates of standard neat human serum samples for both enzymes. **A.** In the presence of trypsin, each peptide's normalized response is reported in replicates of neat human serum. **B.** In the presence of trypsin/Lys-C mixture enzyme, each peptide's normalized response is reported in replicates of neat human serum. **C.** CV values of peptides digested with trypsin **D.** CV values of peptides digested with trypsin/Lys-C mixture.

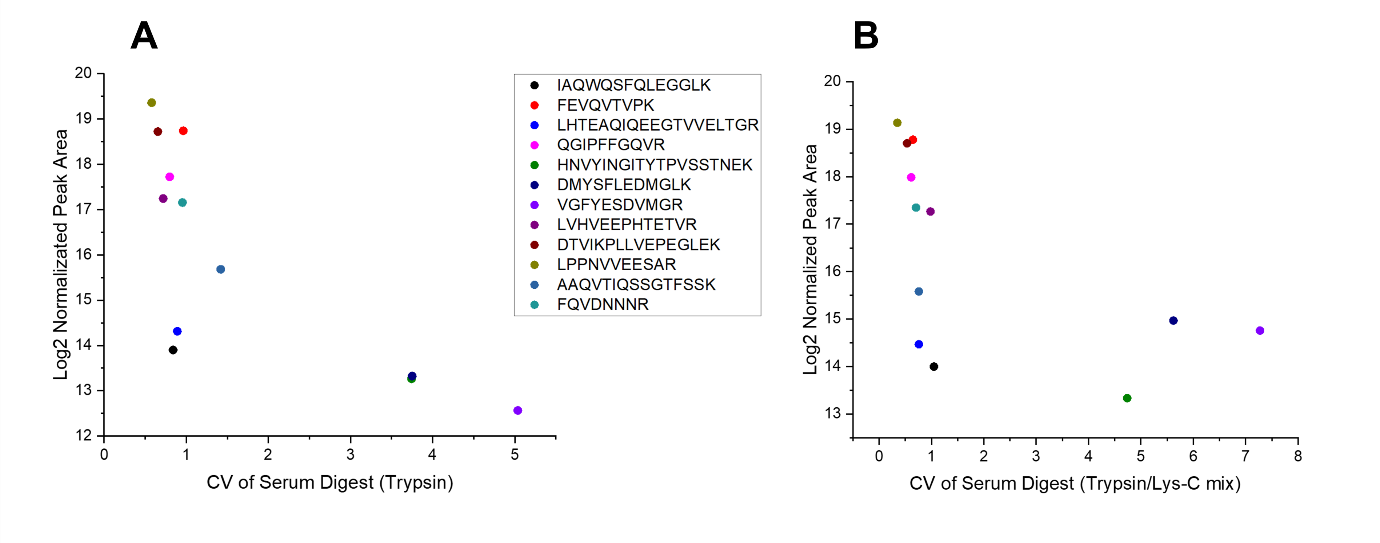

**Figure S4**. The Log2 normalized peak areas versus CV values of neat human serum samples. **A.** in trypsin **B.** in trypsin/Lys-C mixture

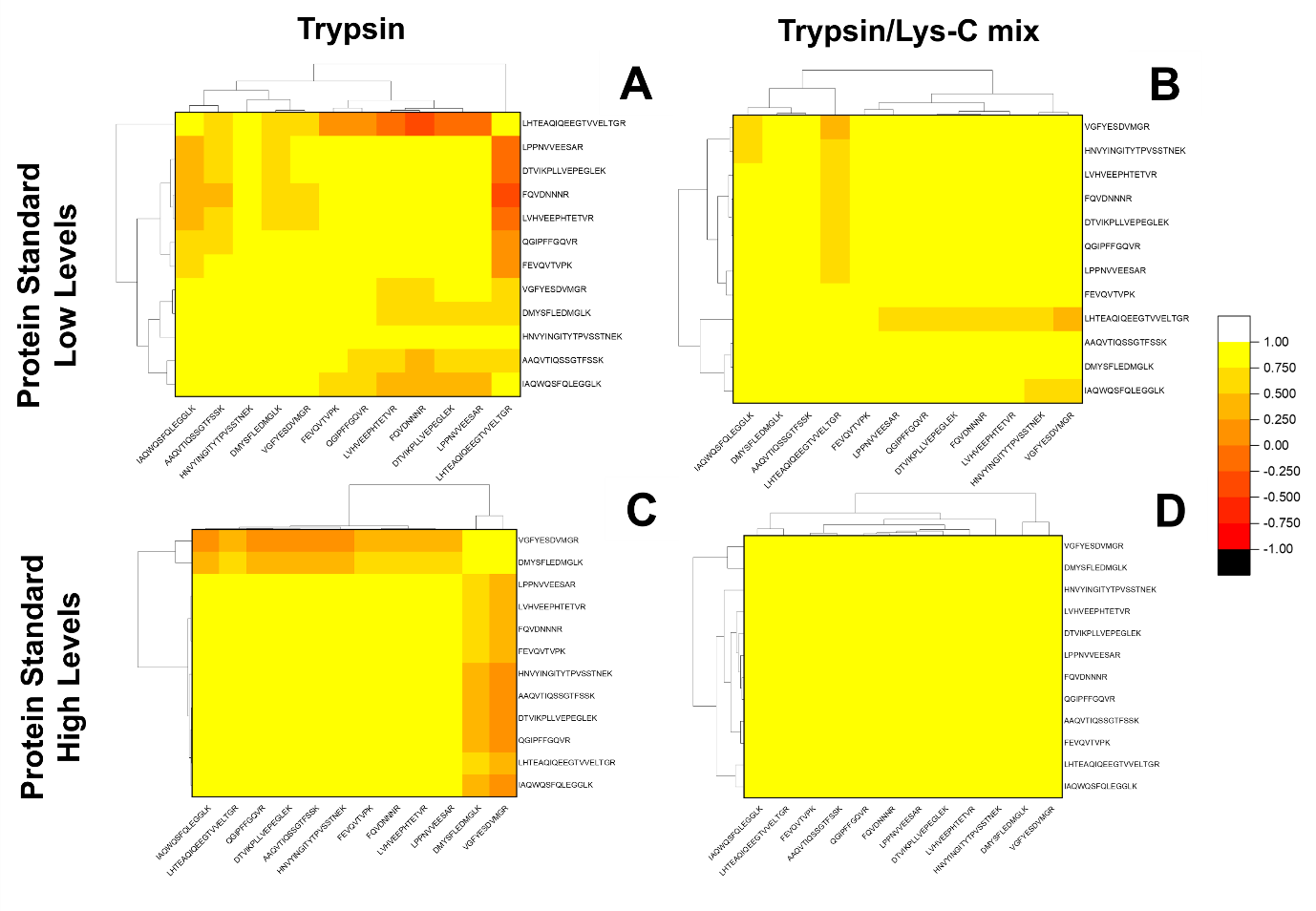

**Figure S5.** A heat map with a dendrogram plot demonstrating the association of A2MG unique peptides in protein standard samples presents different enzymes. **A.** Correlation in low concentration protein standard samples in the presence of trypsin. **B.** Correlation in low-concentration protein standard samples with trypsin/Lys-C mixture. **C.** The correlation in high concentration protein standards presence of trypsin. **D.** Correlation in high-concentration protein standard samples with trypsin/Lys-C mixture.

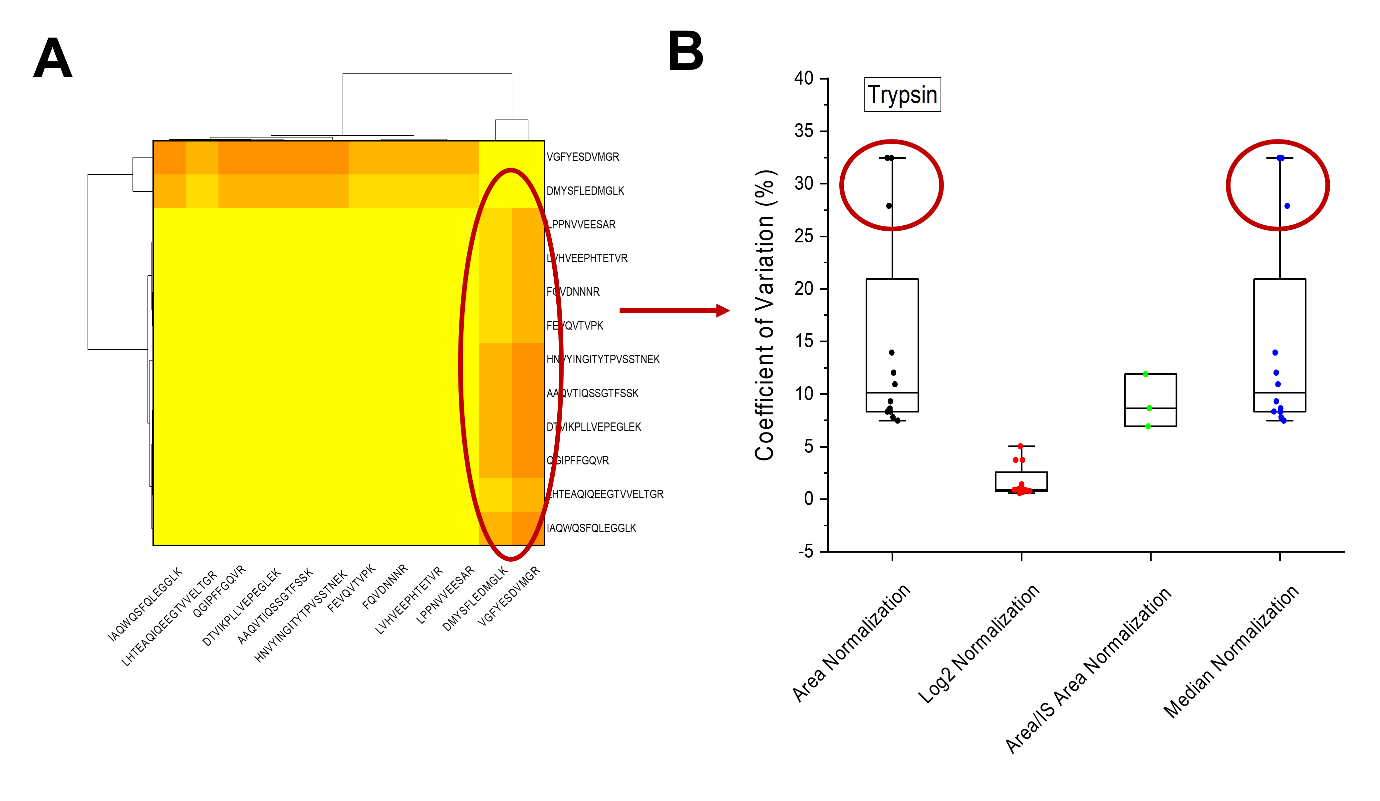

**Figure S6.**The oppositely correlated peptides in **Figure S5C** with the highlighted CV values. **A.** The peptide correlation due to trypsin digestion in **Figure S5C**. The oppositely correlated peptides are highlighted. **B.** The CV values of these peptides are highlighted.

**Table S7.** List of A2MG protein’s unique peptides shared to both matrices. *The peptide has not been included due to low ionization during MS analysis **The synthetic unique peptides were purchased.

|  | **Peptide Sequence** |
| --- | --- |
| 1 | HNVYINGITYTPVSSTNEK |
| 2 | DMYSFLEDMGLK** |
| 3 | DTVIKPLLVEPEGLEK |
| 4 | LPPNVVEESAR** |
| 5 | ALLAYAFALAGNQDK* |
| 6 | AAQVTIQSSGTFSSK |
